## Supplementary Fig. for "Glucomannan engineering highlights the role of galactosyl modification in fine-tuning cellulose-glucomannan interaction in Arabidopsis cell walls"

Contents

- Supplementary Fig. 1 A phylogenetic tree of the GT34 family.
- Supplementary Fig. 2 MAGT expression in *N. benthamiana* leaves and activity on the other substrates.
- Supplementary Fig. 3 Determination of oligosaccharides structures found *in vitro*.
- Supplementary Fig. 4 Comparison of Alphafold models of MAGTs.
- Supplementary Fig. 5 Complimentation of *magt1* mutant in two other individual lines.
- Supplementary Fig. 6 Expression of MAGT has no effect on the backbone structure of AcGGM or other polysaccharides.
- Supplementary Fig. 7 Determination of oligosaccharide structures found in *pIRX3::MAGT* lines.
- Supplementary Fig. 8 Majority of AcGGM in *pIRX3::MAGT* lines remains extractable in the KOH fraction.
- Supplementary Fig. 9 AcGGM is an immobile polymer in wild-type Arabidopsis.

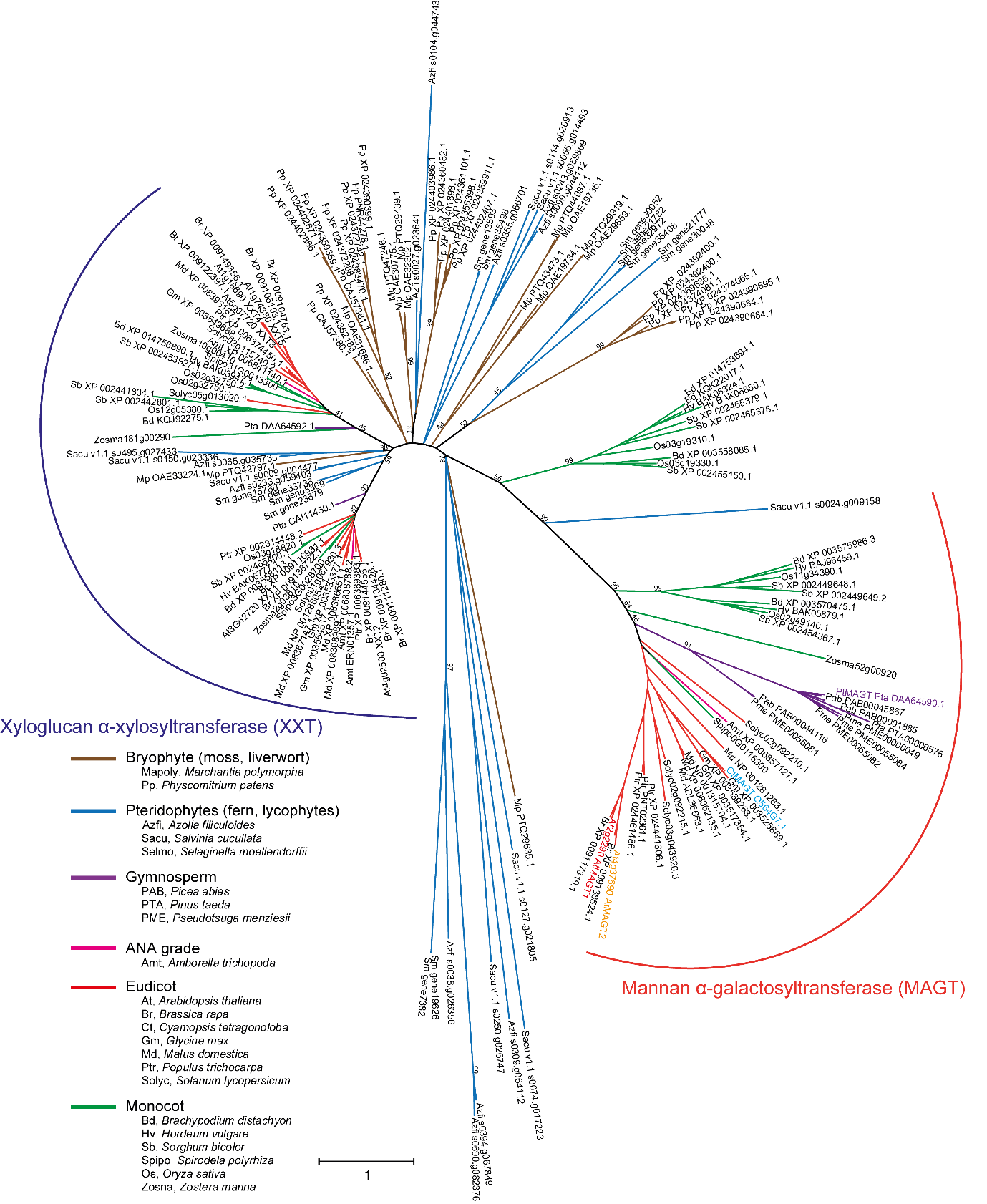

**Supplementary Fig. 1 A phylogenetic tree of the GT34 family.** Bootstrap values at major branching points are shown. Protein alignment was conducted by MUSCLE algorithm with 175 protein sequences collected by BLAST search from NCBI (<https://www.ncbi.nlm.nih.gov/>) and PLAZA (<https://bioinformatics.psb.ugent.be/plaza/>), using AtMAGT1 sequences as a query, and the tree was built by the maximum likelihood algorithms in MEGA X software^1^. Af, *Azolla filiculoides*; Amt, *Amborella trichopoda*; At, *Arabidopsis thaliana*; Bd, *Brachypodium distachyon*; Br, *Brassica rapa*; Ct, *Cyamopsis tetragonoloba*; Gm, *Glycine max*; Hv, *Hordeum vulgare*; Md, *Malus domestica*; Mp, *Marchantia polymorpha*; Os, *Oryza sativa*; Pa, *Picea abies*; Pm, *Pseudotsuga menziesii*; Pp, *Physcomitrium patens*; Pta, *Pinus taeda*; Ptr, *Populus* *trichocarpa*; Sb, *Sorghum bicolor*; Sc, *Salvinia cucullata*; Sl, *Solanum lycopersicum*; Sm, *Selaginella moellendorffii*; Sp, *Spirodela polyrhiza*; Zm, *Zostera marina*. The protein sequences used here are listed in Supplementary Data 2.

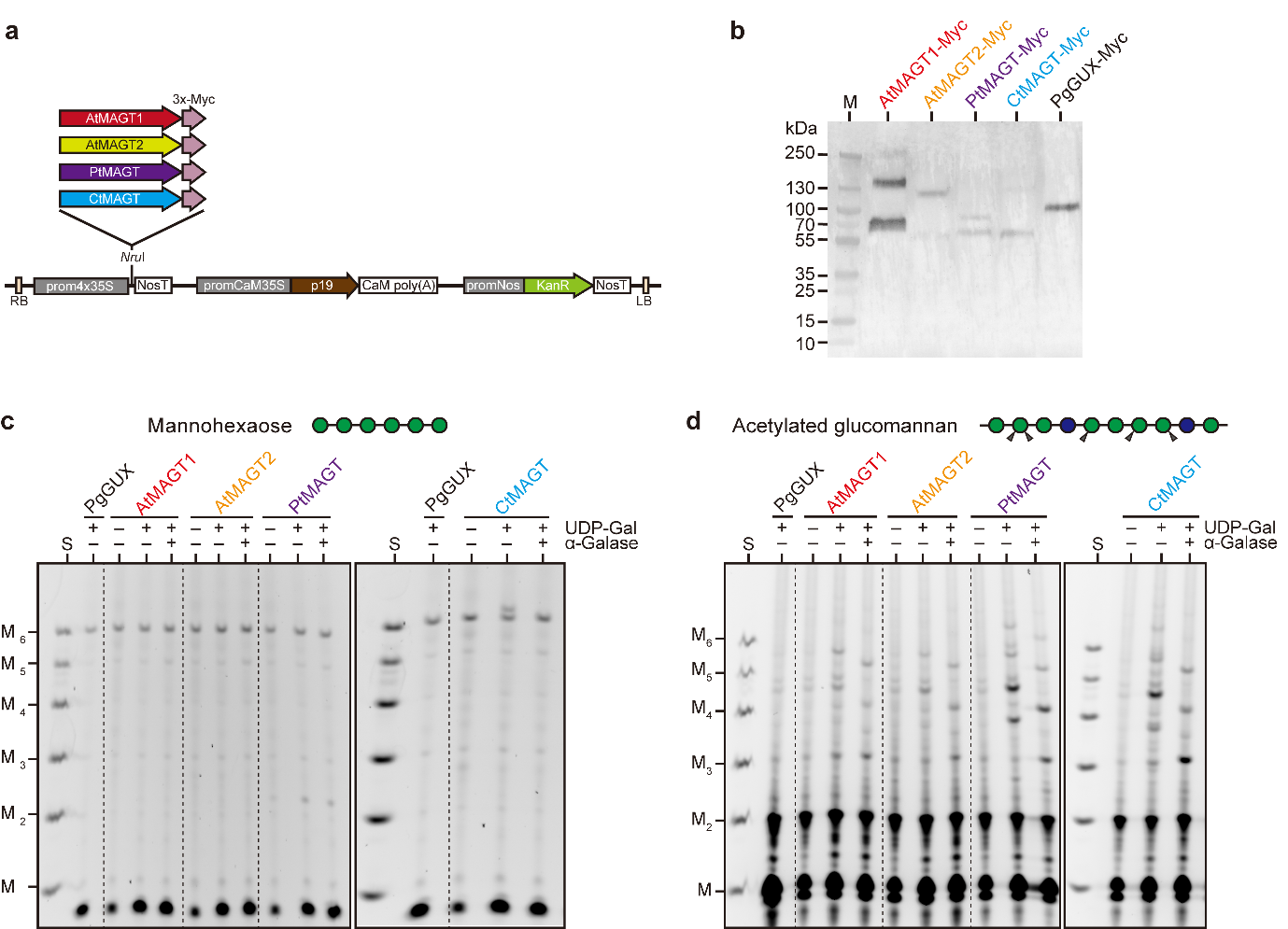

**Supplementary Fig. 2 MAGT expression in *N. benthamiana* leaves and activity on the other substrates. a** A vector map of MAGT expression construct for the transient expression in *N. benthamiana* leaves. Myc-fusions of MAGTs were cloned at *Nru*I site of the pEAQ-HT vector. 4x35S promoter, (prom4x35S); NosT, nos terminator; p19, p19 silencing suppressor; CaM poly(A), cauliflower mosaic virus 35S terminator; promNos, nos promoter; KanR, kanamycin-resistant gene. **b** immune blot of MAGT proteins expressed in *N. benthamiana*. Five μg of total protein of microsomal membrane fraction was used. The anti-Myc polyclonal antibody was used to detect the MAGTs. Image is a composite of colourimetric and chemiluminescent images. M, protein ladder. **c** *In vitro* activity of MAGTs towards mannohexaose. **d** *In vitro* activity of MAGTs towards acetylated glucomannan from pine wood. Galactosylation was confirmed by α-galactosidase (α-Gal) treatment. S, standards of Man and mannooligosaccharides with DP 2-6.

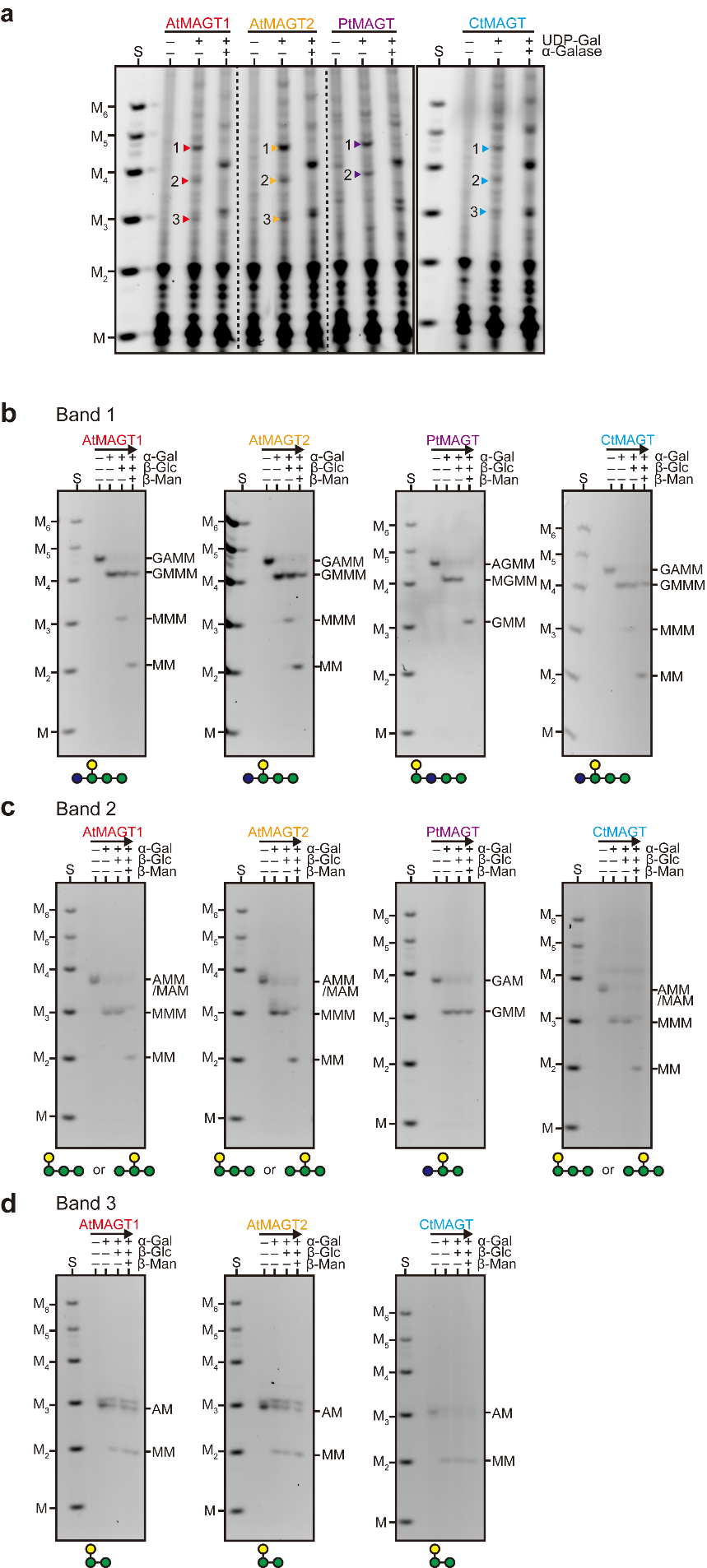

**Supplementary Fig. 3 Determination of oligosaccharide structures found *in vitro*.** (**a)** Selection of bands for the extraction. Bands were numbered from the top. Band 1 (**b**), Band 2 (**c**), and Band 3 (**d**) were extracted from the gel and subjected to further analysis by sequential digestion in the order of α-galactosidase (α-Gal), β-glucosidase (β-Glc), and β-mannosidase (β-Man). The determined structures were illustrated below the gels. GMM and MM were not further digested by the glycoside hydrolases due to the presence of ANTS at the reducing end of the molecules. Given that the β-glucosidase/β-mannosidase digestion prior to the analysis, the position of the Gal side chain should be at the first or second residue from the non-reducing end (e.g. GMAM would have been digested into MAM). For AMM/MAM, the position of Gal substitution at either first or second from the non-reducing end was deduced as they were tolerant with β-Man treatment. S, standards of Man and mannooligosaccharides with D.P. 2-6.

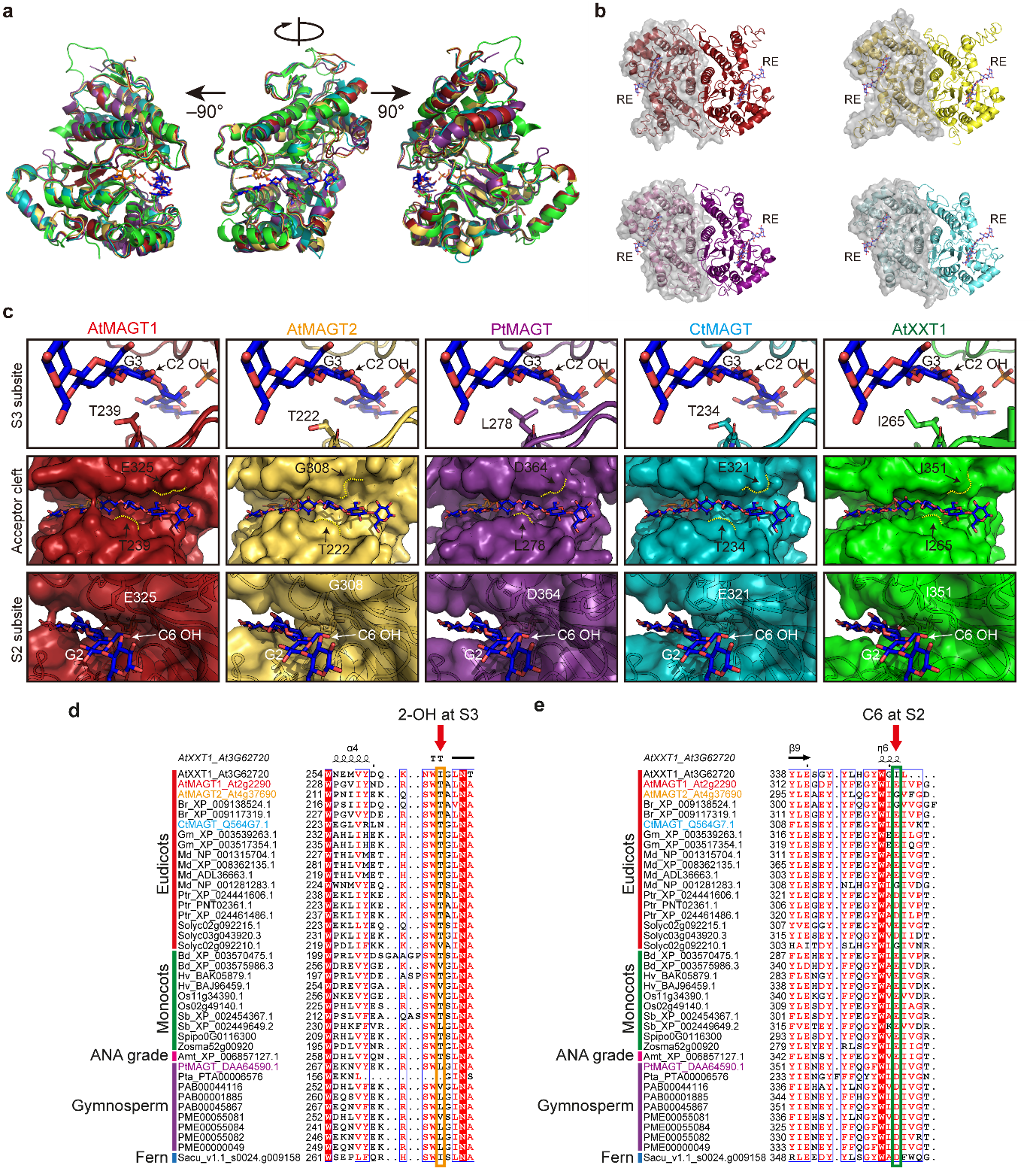

**Supplementary Fig. 4 Comparison of Alphafold models of MAGTs.** **a** Whole view of the model catalytic domain of MAGTs predicted by ColabFold^2^. The protein structures are overlayed with XXT1 crystal structure, UDP and cellohexaose (PBD: 6BSW)^3^. AtMAGT1, AtMAGT2, PtMAGT, CtMAGT, and AtXXT1 were shown in red, yellow, purple, cyan and green, respectively. UDP and Cellohexaose are shown in orange and blue, respectively. **b Homo**dimer form of MAGTs. RE indicates the reducing end of acceptor. **c** Close-up view of subsite 3, where the third glucosyl residue from the reducing end of cellohexaose binds. PtMAGT has a non-polar residue beneath the acceptor whereas the other MAGTs have threonine, which potentially forms a hydrogen bond with C2 hydroxyl of Man that is in an axial position in comparison to an equatorial position in Glc. **d** Close-up view of subsite 2. The glycine of AtMAGT2 makes a pocket for the galactosyl branch on the acceptor. Dotted lines show the surface of the acceptor pocket. Images were taken by Pymol software. **e** Conserved non-polar residues at subsite 3 beneath the acceptor in gymnosperms. **f** Glycine at subsite 2 found in multiple eudicots. Multiple alignments of MAGTs from various plant species were generated by MUSCLE in MEGA X^1^. The alignment was depicted by ESPript 3.0 web (https://espript.ibcp.fr)^4^.

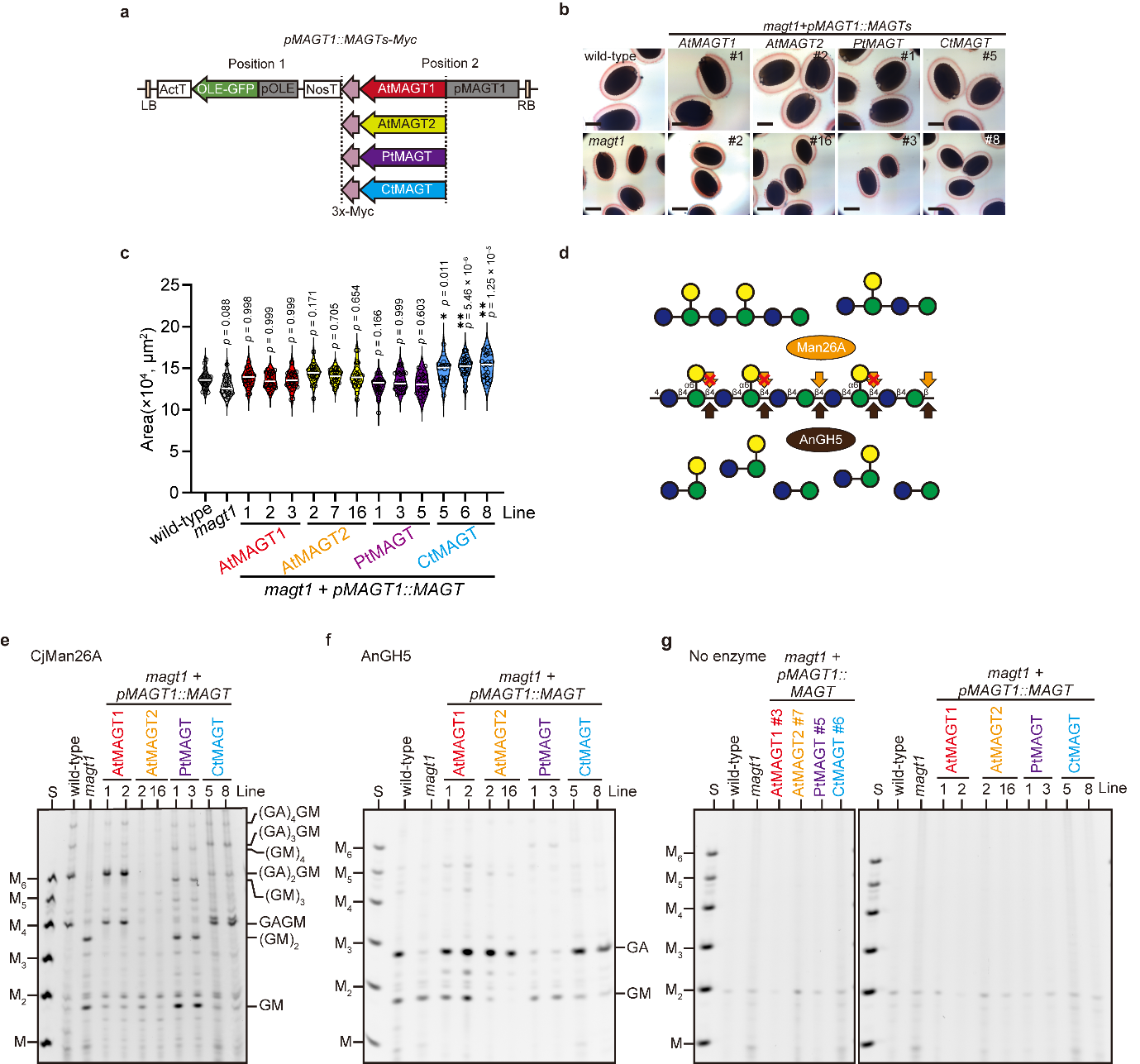

**Supplementary Fig. 5 Comprimentation of *magt1* mutant in two other individual lines. a** Vector map of the construct used for complementation of *magt1* mutant. Position 1 contains a selection marker of OLE-GFP and position 2 has MAGT genes under the promoter of *AtMAGT1*. **b** Mucilage capsule of the other complementation lines. **c** The area of seed measured. Open circles indicate individual measurements; the white lines represent the median of the group. One-way ANOVA (two-tailed) indicated a significant effect of genotype on seed area (*p* = 1.00 $\times$ 10^-20^; *F_13,335_* = 11.61). Results of post hoc multiple comparisons (Dunnet’s method compared with wild-type; wild-type, *n* = 26; *magt1*, *n* = 25; *AtMAGT1#1*, *n* = 24; *AtMAGT1#2*, *n* = 25; *AtMAGT1#3*, *n* = 29; *AtMAGT2#2*, *n* = 25; *AtMAGT2#7*, *n* = 25; *AtMAGT2#16*, *n* = 25; *PtMAGT#1*, *n* = 27; *PtMAGT#3*, *n* = 27; *PtMAGT#5*, *n* = 24; *CtMAGT#5*, *n* = 20; *CtMAG#6*, *n* = 26; *CtMAGT#8*, *n* = 21) are indicated by asterisks (*, *p* < 0.05; **, *p* < 0.01) with *p* values. **d** Different substrate recognition by two mannanases, CjMan26A and AnGH5. The subsite -1 of AnGH5 can be accommodated by galactosylated mannosyl residues, whereas CjMan26A cannot. CjMan26A (**e**) and AnGH5 (**f**) digestion profile of mucilage glucomannan from the other lines of complemented lines. S, standards of Man and mannooligosaccharides with D.P. 2-6. **g** Control for undigested materials.

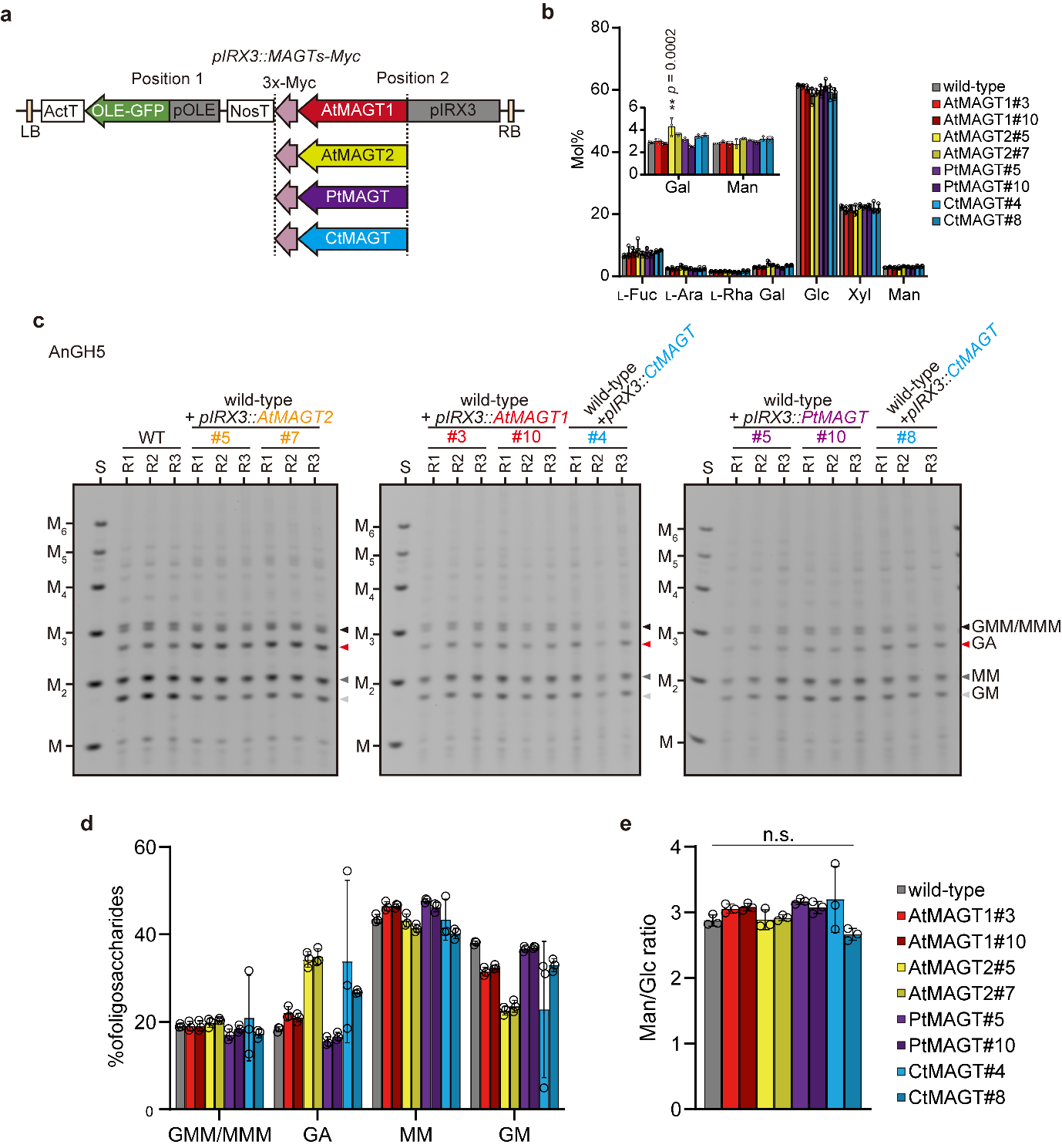

**Supplementary Fig. 6 Expression of MAGT has no effect on the backbone structure of glucomannan nor other polysaccharides. a** Vector map of the construct used for glucomannan engineering. Position 1 contains a selection marker of OLE-GFP and position 2 has MAGT genes under the promoter of *IRX3*. **b** Monosaccharides composition analysis of total cell wall materials from the bottom part of the stem. AIR was hydrolysed by sulphuric acid and monosaccharides were measured by HPAEC-PAD. Data are mean values with standard deviations of three biological replicates. One-way ANOVA (two-tailed) was performed on each monosaccharide. Monosaccharides shown significant difference was further analysed by Dunnett’s multiple comparison test (**, *p* < 0.01). F values and P values were provided in Supplementary Data 5. **c** AnGH5 digestion profile of KOH fraction obtained from *pIRX3::MAGT* lines was used to calculate Gal/Man ratio in Fig. 3. Bands used for calculation were annotated with arrowheads. S, standards of Man and mannooligosaccharides with D.P. 2-6. **d** Proportion of oligosaccharides based on the band intensity. **e** Man/Glc ratio. Data are mean values and standard deviation of three biological replicates. No significant difference detected by one-way ANOVA (two-tailed; *p* = 0.0528; *F_8,18_* = 2.472) confirmed that the glucomannan backbone structure was not affected in *pIRX3::MAGT*.

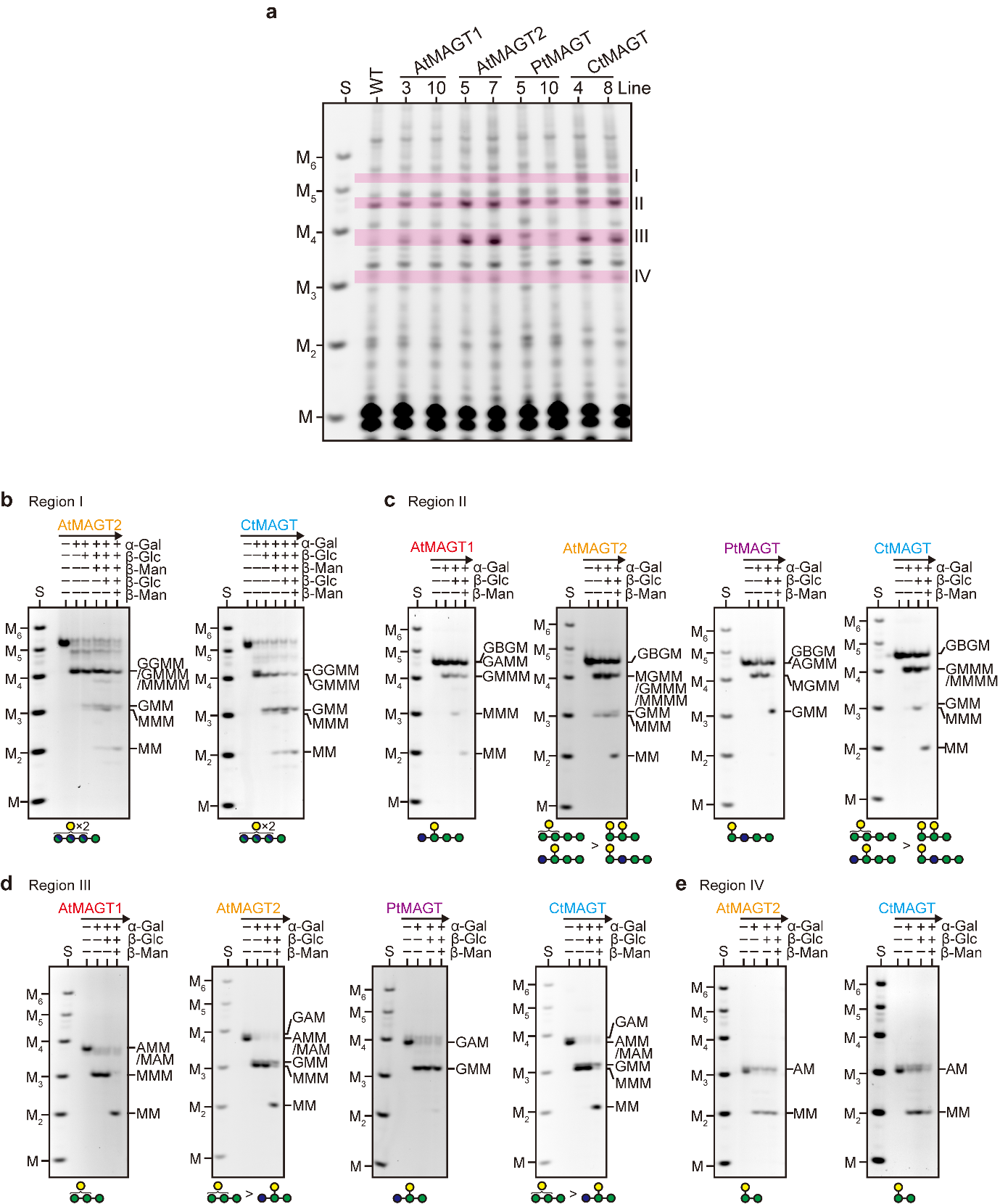

**Supplementary Fig. 7 Determination of oligosaccharides structures found in *pIRX3::MAGT* lines.** **a** Galactosylated products found in two independent *pIRX3::MAGT* lines. Unique bands in the region I (**b**), II (**c**), III (**d**), and IV (**e**) were extracted for further analysis by sequential digestion with α-galactosidase (α-Gal), β-glucosidase (β-Glc), and β-mannosidase (β-Man). S, standards of Man and mannooligosaccharides with D.P. 2-6. The determined structures were illustrated below the gels. GMM and MM were not further digested by the glycoside hydrolases due to the presence of ANTS at the reducing end of the molecules. The oligosaccharides in the region I had several backbone structures, such as MGMM, GMMM, and MMMM, with two Gal modifications. The oligosaccharides in region II of *pIRX3::AtMAGT2* showed MMM after α-Gal digestion, indicating the presence of AAM structure. It should be noted that the tolerant bands after sequential digestion in region II are likely to be GBGM (where B is a β-Gal-1,2-α-Gal-1,6-Man unit) derived from β-GGM in primary cell walls^5^. For AMM/MAM in region III, the position of Gal substitution at either first or second from the non-reducing end was deduced as they were tolerant with β-Man treatment.

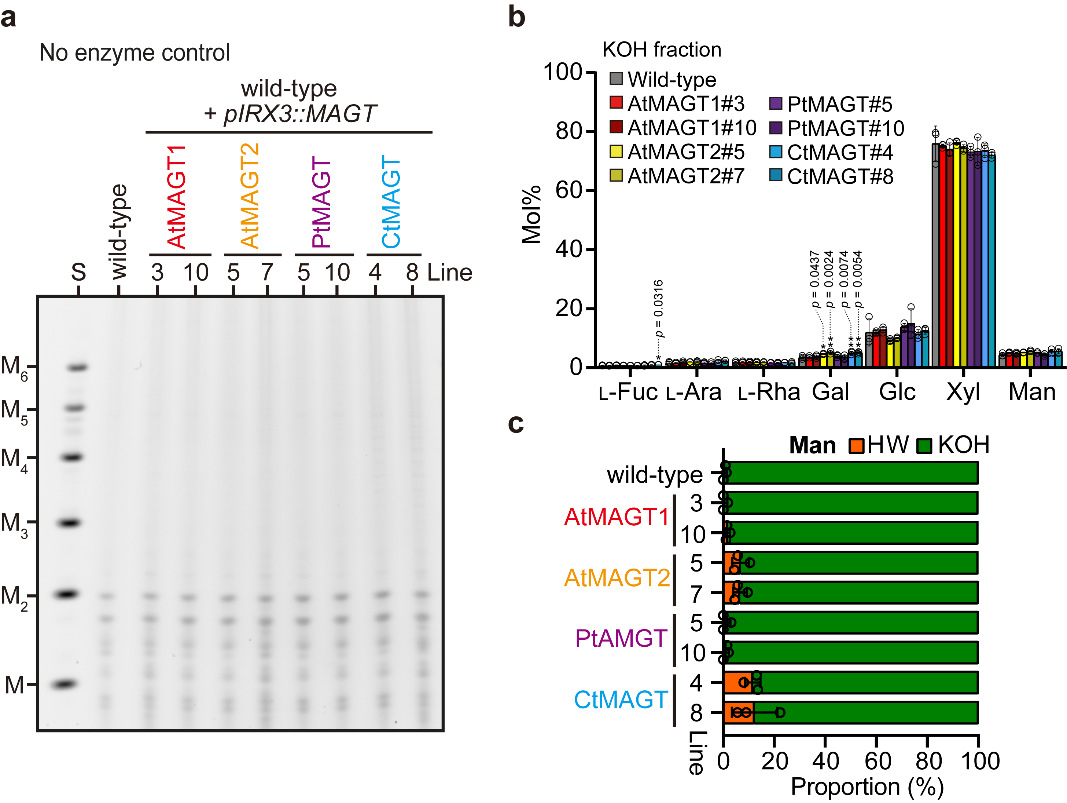

**Supplementary Fig. 8 Majority of glucomannan in *pIRX3::MAGT* lines remains extractable in the KOH fraction. a** Control of undigested materials of hot water (HW) fraction. S, standards of Man and mannooligosaccharides with D.P. 2-6. **b** Monosaccharides composition analysis of KOH fraction after the sequential extraction by HW and ammonium oxalate. Data are mean values with standard deviations of three biological replicates. Open circles indicate individual measurements. One-way ANOVA (two-tailed) was performed on each monosaccharide. Monosaccharides shown significant difference was further analysed by Dunnett’s multiple comparison test (*, *p* < 0.05; **, *p* < 0.01). F values and P values were provided in Supplementary Data 6. **c** Proportion of fractions where Man extracted. Data are mean values with standard deviations of three biological replicates. Open circles indicate individual measurements.

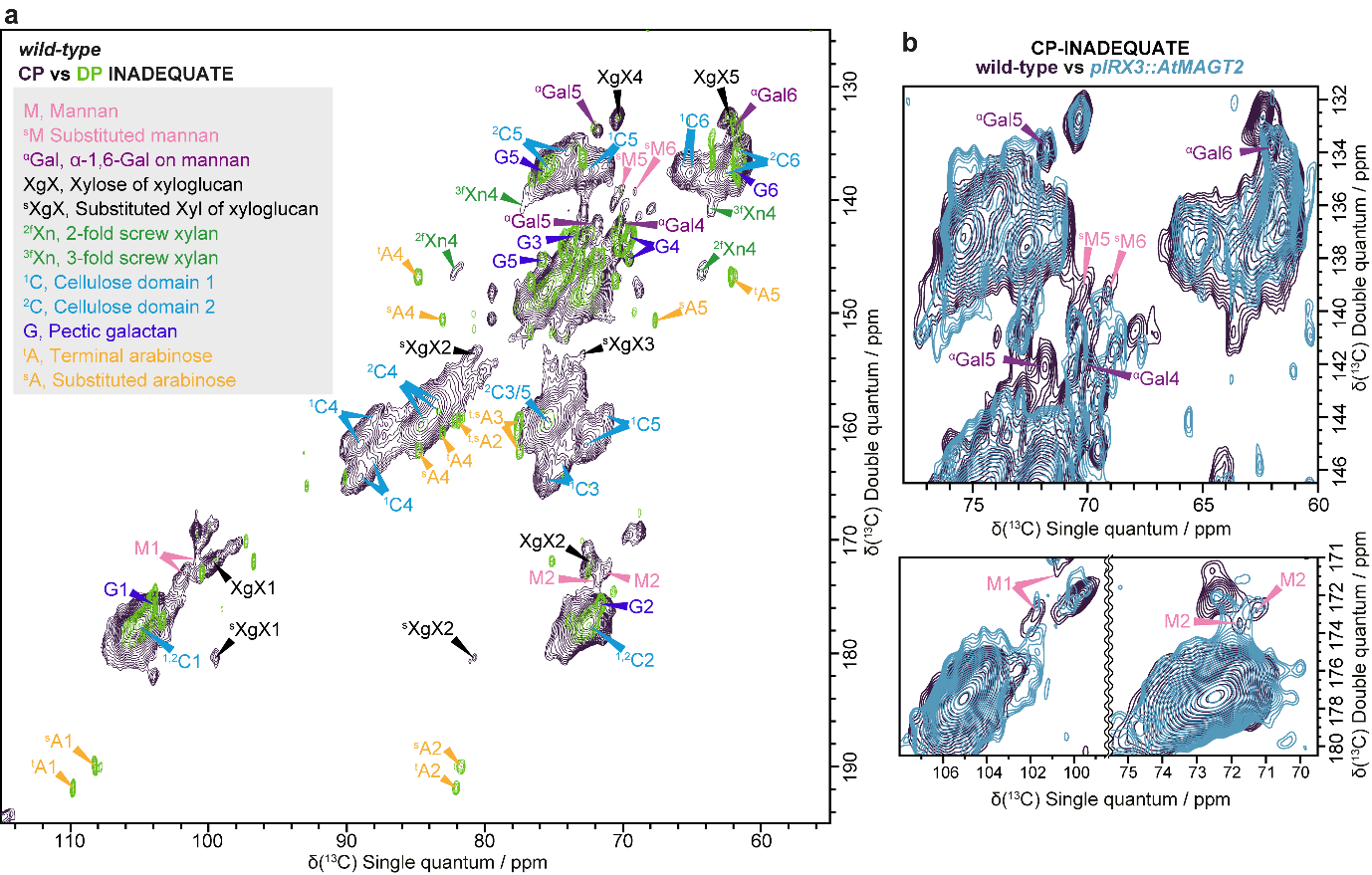

**Supplementary Fig. 9 Glucomannan is an immobile polymer in wild-type.** Solid-state NMR spectra of Arabidopsis wild-type inflorescence stem. **a** Overlay of ^13^C CP- and DP-refocused INADEQUATE MAS ssNMR spectra of wild-type. AcGGM peaks are labelled: Man (M), α-Gal (^α^Gal). Cellulose (domain 1, ^1^C; domain 2, ^2^C), xylan (two-fold screw, ^2f^Xn; three-fold screw, ^3f^Xn), xylose of xyloglucan (XgX), arabinose (A), and pectic galactan are also labelled. Most AcGGM peaks were found in the CP spectrum, that is, AcGGM is immobile in the cell walls of wild-type, suggesting that it binds to cellulose. **b** Comparison of C1, C2, and C5/6 regions of the CP spectra between wild-type and *pIRX3::AtMAGT2*. Much less intensity of AcGGM peaks in CP spectra of *pIRX3::AtMAGT2* compared to wild-type. Spectra were acquired at a ^13^C Larmor frequency of 213.8 MHz and a MAS frequency of 12.5 kHz. The spin-echo duration used was 2.24 ms. Chemical shifts of the annotated peaks are listed in Supplementary Data 1.
